# Black flying foxes in Australia harbor novel *Borrelia* lineages adjacent to zoonotic clades

**DOI:** 10.1101/2024.12.20.629695

**Authors:** Taylor B. Verrett, Caylee A. Falvo, Evelyn Benson, Devin N. Jones-Slobodian, Daniel E. Crowley, Adrienne S. Dale, Tamika J. Lunn, Manuel Ruiz-Aravena, Bat One Health, Clifton D. McKee, Kerry L. Clark, Alison J. Peel, Raina K. Plowright, Daniel J. Becker

## Abstract

We explore the role of Australian black flying foxes (*Pteropus alecto*) as reservoir hosts of potentially zoonotic *Borrelia* bacteria. Across six sites, 2% of 840 bats were infected with one of two novel *Borrelia* haplotypes. Phylogenetic reconstruction indicated these infections are distinct from Lyme or relapsing fever clades.

## Introduction

Bacteria in the genus *Borrelia* are causative agents of two diseases that represent significant public health burdens: Lyme borreliosis and relapsing fever. Lyme borreliosis is the most frequently reported vector-borne infection in the northern hemisphere, where it is vectored by ixodid ticks (Khatchikian et al. 2015). By contrast, relapsing fever is globally distributed, vectored predominantly by argasid ticks, and is attributed annually to thousands of cases of febrile illness in humans (Jakab et al. 2022).

Lyme borreliosis and relapsing fever lay the basis for two monophyletic *Borrelia* clades: the *B. burgdorferi* sensu lato (*Bb*sl) complex and the relapsing fever (RF) group. *Borrelia* from both groups are harbored by a broad range of vertebrate hosts, including birds, reptiles, and mammals. Identifying reservoir hosts and clarifying their role in propagating *Borrelia* is critical for monitoring and mitigating spillover risk. For example, migratory birds can contribute to the long-distance dispersal of *Bb*sl by transporting millions of ticks within and across continents (Ogden et al. 2008).

Bats may play an underrecognized role in the dispersal and/or enzootic maintenance of *Borrelia* bacteria. Bats are volant mammals that have been known to harbor borrelial spirochaetes for over a century (Nicolle and Comte 1906), and recent surveys have linked borreliae from both the *Bb*sl and RF groups to chiropteran hosts (Colunga-Salas et al. 2021; Muñoz-Leal et al. 2021). There is also evidence that bat-associated *Borrelia* infections can be zoonotic, as *Borrelia* lineages linked to probable bat reservoirs have been implicated in febrile illness in humans (Qiu et al. 2019). The expansion of *Borrelia* research in chiropteran hosts could therefore inform current and future welfare of both bat and human populations.

Australian flying foxes (*Pteropus* spp., family Pteropodidae) represent a group that is highly prominent at the human–wildlife interface and therefore are an important target for *Borrelia* surveillance. The increasing propensity for flying foxes to roost in or near human settlements could heighten the overlap of humans and bat ectoparasites, opening a potential *Borrelia* spillover pathway (Eby et al. 2023; Szentiványi et al. 2024). While black flying foxes (*Pteropus alecto*) have been the subject of extensive research as reservoir hosts of Hendra virus, relatively little study has been devoted to their bacterial communities, including those that could be pathogenic (Szentivanyi et al. 2023).

### The Study

Here, we assess the presence, diversity, and phylogenetic placement of *Borrelia* spp. associated with black flying foxes in Australia. As part of ongoing research on the ecology of Hendra virus, we collected blood samples from 840 black flying foxes across six sites in southern Queensland and northern New South Wales between May 2018 and September 2020 (Figure 1). Bats were captured with mist nets, anesthetized with isoflurane, and sampled for blood preserved on Whatman FTA cards until further processing (Hansen et al. 2022). Field protocols were approved by the Montana State University Institutional Animal Care and Use Committee (201750) and Griffith University Animal Ethics Committee (ENV/10/16/AEC and ENV/07/20/AEC). We used Qiagen QIAamp DNA Investigator Kits to extract genomic DNA from blood samples, following the manufacturer’s instructions. To determine the presence of *Borrelia* spp., we used PCRs targeting the 16S rRNA gene, flagellin (*flaB*) gene, and 16S–23S rRNA intergenic spacer (IGS). Primers and annealing conditions are provided in Table S1. PCR products of the expected range for positives were purified with the Promega Wizard SV Gel and PCR Clean-Up System and Sanger sequenced in both directions at Eurofins Genomics. We report prevalence data for IGS, but this gene was not included in phylogenetic analyses due to insufficient reference sequences in GenBank. Our *Borrelia* spp. sequences from black flying foxes were edited and trimmed manually and then aligned with reference sequences in GenBank using the MUSCLE algorithm via Geneious 2024.0.5. We constructed single-marker phylogenies using both Bayesian and maximum likelihood (ML) methods. For Bayesian analyses, we used MrBayes 3.2 with 10 million Markov chain Monte Carlo generations and the default burn-in of 25% (Ronquist et al. 2012). For ML analyses, we used RAxML 8.0 with a starting tree obtained by searching for the best-scoring ML tree in a single run and calculating branch support with 1000 rapid bootstrap replicates (Stamatakis 2014). All trees used a GTR+I+G nucleotide substitution model (Abadi et al. 2019).

**Figure 1.**
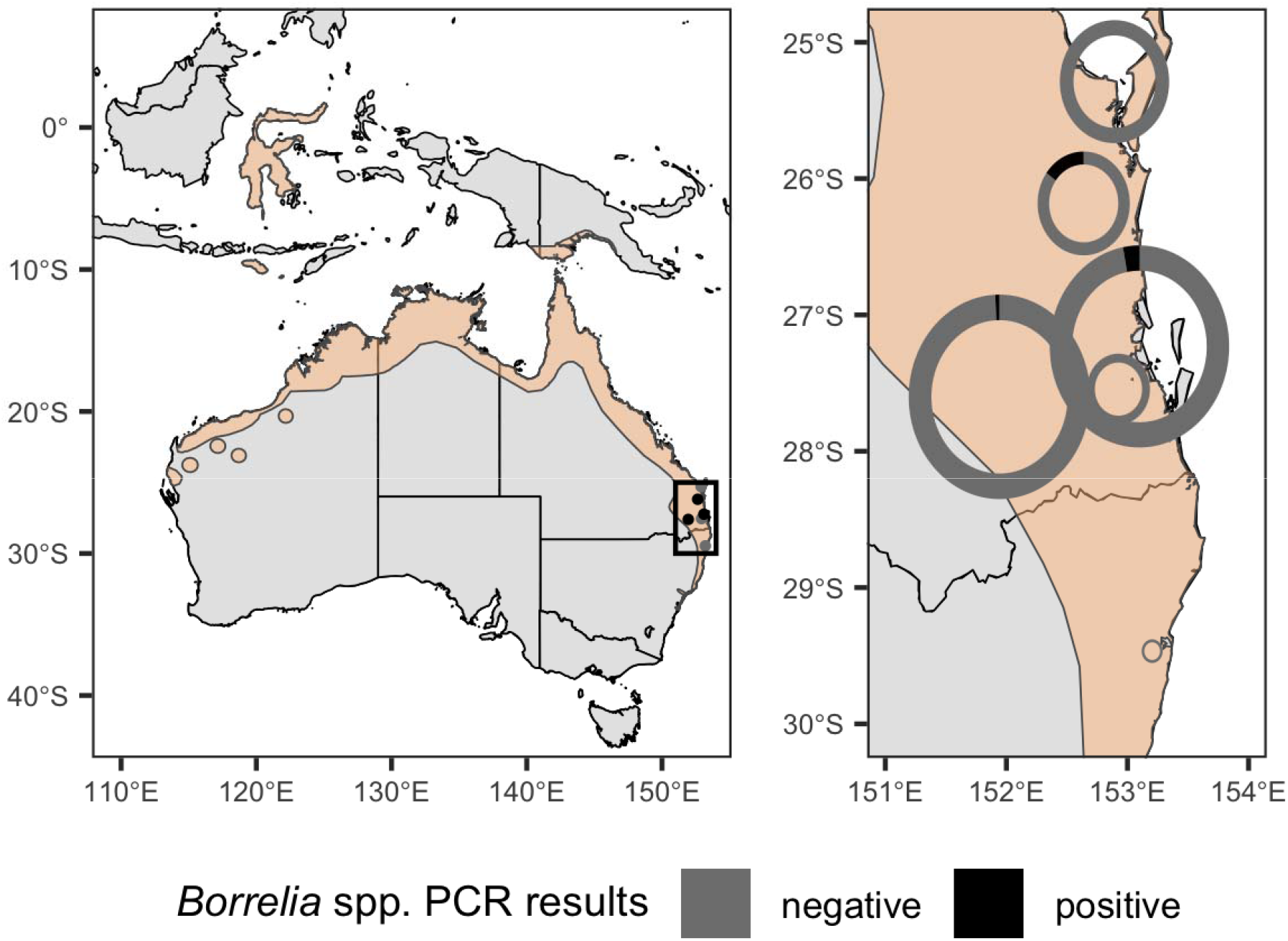
Map of sampling sites in eastern Australia (Queensland and New South Wales) in relation to the *Pteropus alecto* geographic distribution (in orange) as defined by the International Union for Conservation of Nature. Points are colored by the presence or absence of *Borrelia* spp. infections in a given site. The inset displays donut plots with the fraction of PCR-positive samples in black; points are scaled by log-transformed sample size.

Across our approximately two-year study period and six roosts, 2% of black flying foxes tested positive via PCR for *Borrelia* spp. (17/840 samples, 95% CI: 1.3–3.2%; Table 1). Roost-specific infection prevalence ranged from 0% (three of the six roosts) to 15% (3/20 samples, 95% CI: 5.2–36.0%) in Gympie, Queensland. The roosts with the greatest sampling effort (Toowoomba and Redcliffe in Queensland, *n* = 402 and 374) had 0.7% and 2.9% of bats infected, respectively (Table S2). Most infected bats (13/17) yielded usable sequence data from either 16S rRNA or *flaB* gene targets, which represented two haplotypes (p-distances = 8.2% for *flaB* and 1.3% for 16S rRNA). Topologies were similar across tree-building methods (Supplemental Figures S1 and S2) but somewhat discordant between gene targets. The *flaB* phylogeny grouped black flying fox *Borrelia* spp. with lineages from Macaregua Cave in Colombia in a clade sister to *Bb*sl (Figure 2), but 16S rRNA sequence data are not presently available from Macaregua Cave. There are no other sequences from Australian bats. The 16S rRNA phylogeny (Figure 3) was also less effective at resolving relationships across the RF and *Bb*sl groups supported in previous analyses using multiple markers (Loh et al. 2017). All sequences included here are available on GenBank through accessions PQ488732–PQ488741 (16S rRNA), PQ492350–PQ492360 (*flaB*), and PQ490736–PQ490746 (16S–23S IGS).

**Table 1.** Metadata on individual bats PCR-positive for *Borrelia* spp., including unique identifier, capture site and population categorization, month and year of sampling, sex, and GenBank accession(s). Bats are considered infected if testing positive for at least one of our three markers (i.e., 16S rRNA gene, *flaB* gene, and 16S–23S rRNA ITS) through PCR.

| Bat ID | Site | Date | Sex | Age | GenBank |
| --- | --- | --- | --- | --- | --- |
| ACGMP001-1 | Gympie | 1/2019 | Female | Adult | PQ492350<sup>†</sup>,<br>PQ490736<sup>‡</sup> |
| ACGMP001-4 | Gympie | 1/2019 | Female | Adult | PQ492351<sup>†</sup>,<br>PQ490737<sup>‡</sup> |
| ACGMP001-5 | Gympie | 1/2019 | Female | Adult | PQ488732*,<br>PQ490738<sup>‡</sup> |
| ACRED001-15 | Redcliffe | 5/2018 | Male | Adult | PQ488733*,<br>PQ492352<sup>†</sup>,<br>PQ490739<sup>‡</sup> |
| ACRED003-23 | Redcliffe | 9/2018 | Male | Subadult | PQ488734*,<br>PQ492353<sup>†</sup>,<br>PQ490740<sup>‡</sup> |
| ACRED004-31 | Redcliffe | 12/2018 | Female | Adult | PQ488735* |
| ACRED006-54 | Redcliffe | 5/2019 | Female | Adult | PQ488736*,<br>PQ492354<sup>†</sup>,<br>PQ490741<sup>‡</sup> |
| ACRED007-51 | Redcliffe | 7/2019 | Female | Adult | PQ488737*,<br>PQ492355<sup>†</sup>,<br>PQ490742<sup>‡</sup> |
| ACRED008-54 | Redcliffe | 9/2019 | Female | Adult | PQ492356<sup>†</sup>,<br>PQ490743<sup>‡</sup> |
| ACRED009-72 | Redcliffe | 12/2019 | Male | Subadult | PQ488738*,<br>PQ492357<sup>†</sup> |
| ACRED010-32 | Redcliffe | 3/2020 | Male | Adult | PQ488739*,<br>PQ492358<sup>†</sup> |
| ACRED010-42 | Redcliffe | 3/2020 | Female | Adult | PQ488740*,<br>PQ490744<sup>‡</sup> |

Table 1 (continued). Metadata on individual bats PCR-positive for *Borrelia* spp., including unique identifier, capture site and population categorization, month and year of sampling, sex, and GenBank accession(s).
|  |  |  |  |  |  |
| --- | --- | --- | --- | --- | --- |
| ACRED011-45 | Redcliffe | 5/2020 | Female | Adult | PQ492359†, PQ490745‡ |
| ACRED011-60 | Redcliffe | 5/2020 | Male | Adult | PQ488741* , PQ492360†, PQ490746‡ |
| ACTOW001-15 | Toowoomba | 6/2018 | Female | Adult | No sequences |
| ACTOW002-30 | Toowoomba | 7/2018 | Female | Subadult | No sequences |
| ACTOW006-21 | Toowoomba | 3/2019 | Female | Subadult | No sequences |

\*16S rRNA
†*flaB*
‡16S–23S rRNA IGS

**Figure 2.**
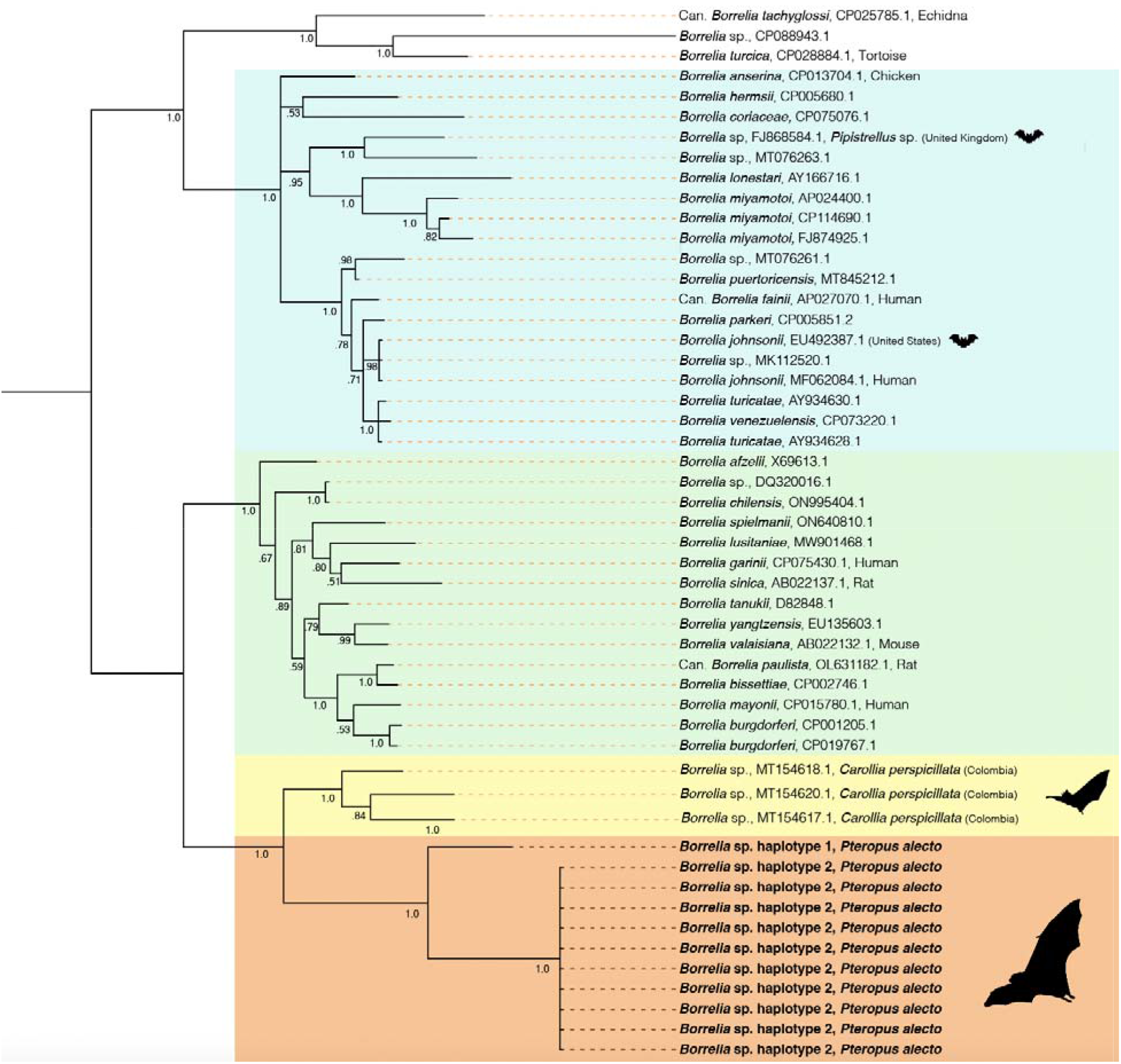
Bayesian phylogenetic tree displaying evolutionary relationships between *Borrelia* spp. using the *flaB* gene. This tree was constructed using a GTR+I+G substitution model and 10 million MCMC generations. Colors represent major *Borrelia* groups and clades of interest: blue = relapsing fever group (RF), green = *Borrelia burgdorferi* sensu lato complex (*Bb*sl), yellow = *Borrelia* spp. from Macaregua Cave, and orange = new *Borrelia* haplotypes from black flying foxes in Australia. Host associations are noted when Genbank lineages were isolated from vertebrates, barring experimental infections. Tips associated with bats or bat ticks are marked with a bat graphic (sourced from Noun Project).

**Figure 3.**
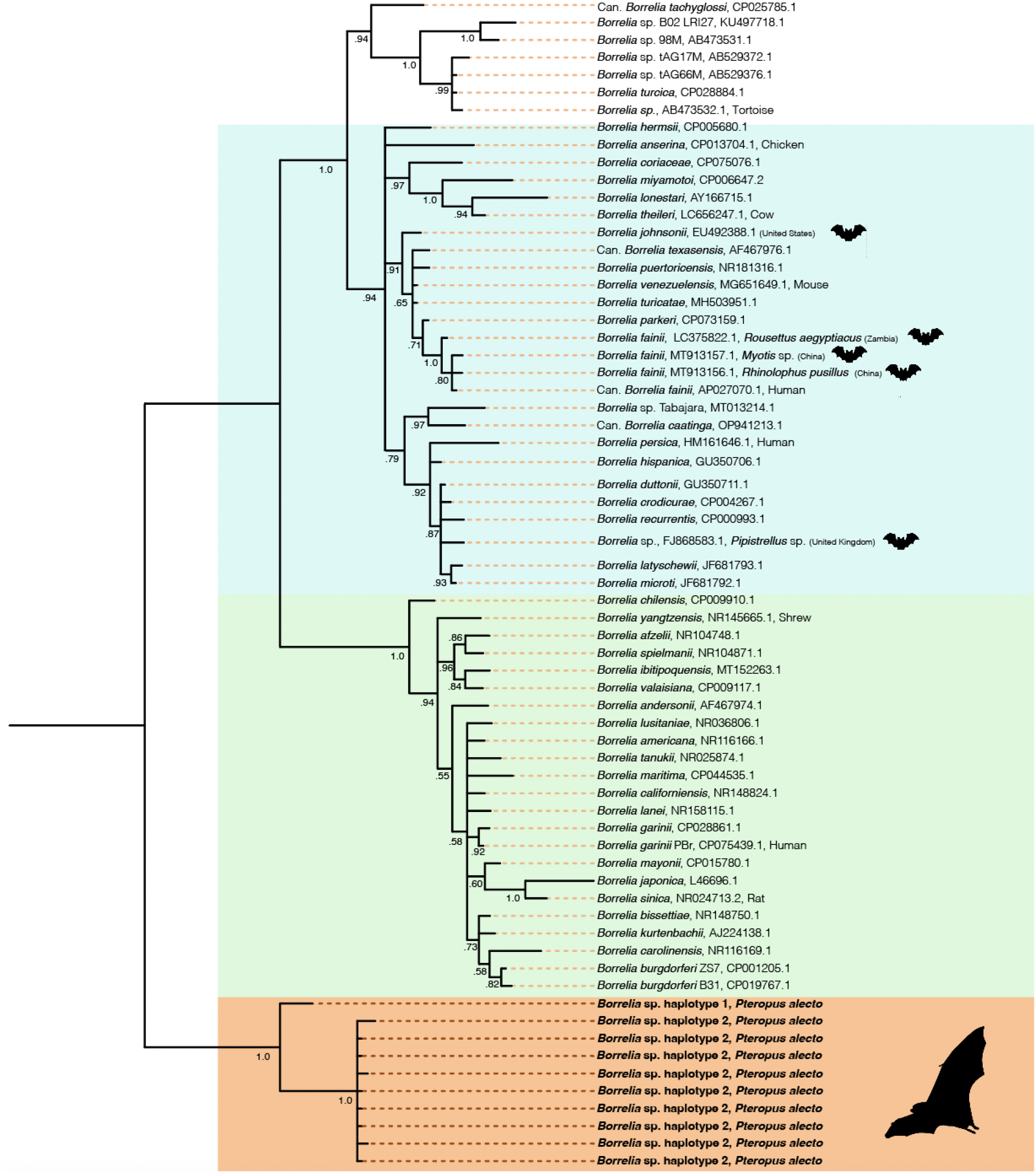
Bayesian phylogenetic tree displaying evolutionary relationships between *Borrelia* spp. using 16S rRNA. This tree was constructed using a GTR+I+G substitution model and 10 million MCMC generations. Colors represent major *Borrelia* groups and clades of interest: blue = relapsing fever group (RF), green = *Borrelia burgdorferi* sensu lato complex (*Bb*sl), and orange = new *Borrelia* haplotypes from black flying foxes in Australia. Host associations are noted when Genbank lineages were isolated from vertebrates, barring experimental infections. Tips associated with bats or bat ticks are marked with a bat graphic (sourced from Noun Project).

## Conclusions

This study represents the first survey of *Borrelia* spp. in Australian bats and corroborates that bats can host *Borrelia* infections evolutionarily distinct from recognized clades, but more closely related to *Bb*sl than RF infections. Sequences from two *Borrelia* spp. lineages were recovered from 11 of 17 infected bats (Figure 2). Haplotype 1 was represented by a single host that shared a roost (Gympie) with a bat infected with haplotype 2, suggesting this variation is unlikely to be structured by geographic isolation. Phylogenetic reconstruction of *flaB* and 16S rRNA suggested *Borrelia* infections from black flying foxes belong to a clade adjacent to existing *Bb*sl and RF groups, and are most closely related to *Borrelia* spp. hosted by phyllostomid bats in Colombia (Muñoz-Leal et al. 2021).

Our results are a base for establishing the presence and phylogenetic placement of *Borrelia* infections in flying foxes, but underscore research gaps in characterizing their zoonotic potential. The virulence of these lineages in flying foxes, for example, is unknown; while bats are tolerant of multiple viruses (Irving et al. 2021), lethal borrelial infections in bats are not unheard of (Evans et al. 2009). Importantly, the arthropod vector remains unidentified, but the host specificity and geographic range of this vector should strongly influence zoonotic risk. Certain ixodid bat ticks, for example, have generalist feeding habits that could position them as an epidemiological link between bats, domestic animals, and humans (Szentiványi et al. 2024). Targeted efforts to sample black flying foxes and their ectoparasites across their range will better inform the zoonotic potential of the novel *Borrelia* lineages described in this study.

## Acknowledgements

We acknowledge the Barunggam, Turrbal, Gubbi Gubbi and Ningy Ningy people, who are the Traditional Custodians of the land upon which this work was conducted. We would like to thank Peggy Eby and Liam McGuire for their ecological insights and contributions to study design as well as Maureen Kessler, Julia van Velden, Ruby Lovelock, Lisa Albino, Ariana Ananda, and Kerryn Parry-Jones for their assistance in the field. We also thank Sara LaTrielle, Isaac Knights, Dian Riseley, and Stella Maris Januario da Silva for project support. Field protocols were approved by the Griffith University Animal Ethics Committee (Certificate: ENV/10/16/AEC and ENV/07/20/AEC), and Montana State University Institutional Animal Care and Use Committee (201750), and research was conducted with a Scientific Purposes Permits from the Queensland Department of Environment and Heritage Protection (WISP17455716 and WA0012532), a permit to Take, Use, Keep or Interfere with Cultural or Natural Resources (Scientific Purpose) from the Department of National Parks, Sport and Racing (WITK18590417), a Scientific License from the New South Wales Parks and Wildlife Service (SL101800) and general and products liability protection permit (GRI 18 GPL), and with permission to undertake research on council and private land. Sample import to Montana State University was authorized by import permit 20200728-2504A.

This work was funded by the DARPA PREEMPT program Cooperative Agreement (D18AC00031) and the U.S. National Science Foundation (DEB-1716698; EF-2133763/ EF-2231624). The content of the information does not necessarily reflect the position or the policy of the U.S. government, and no official endorsement should be inferred. Molecular analyses were also partially supported by the Early Career Grants Programme of the Royal Society of Tropical Medicine and Hygiene.

