## Supplemental figures for "Black flying foxes in Australia harbor novel *Borrelia* lineages adjacent to zoonotic clades"

Supplementary material

**Table S1.** PCR primers and reaction conditions used for *Borrelia* sp. PCR testing.

| **Target Gene** | **Primer Name** | **Primer Orientation** | **5’-3’ sequence** | **Annealing temp (°C)*** | **Cycles** | **Amplified fragment (bp)** | **Reference** |
| --- | --- | --- | --- | --- | --- | --- | --- |
| 16S rRNA | 1A | Forward | CTAACGCTGGCAGTGCGTCTTAAGC | 70-61, 60 | 10 + 40 | ~724 | (Richter et al. 2003) |
|  | 1B | Reverse | AGCGTCAGTCTTGACCCAGAAGTTC |  |  |  |  |
| 16S rRNA | Bf1_24_ | Forward | GCTGGCAGTGCGTCTTAAGCATGC | 72-63, 62 | 10 + 40 | ~1,395 | Modified from (Raoult et al. 1998) |
|  | Br1_24_ | Reverse | GCTTCGGGTATCCTCAACTCGGGT |  |  |  |  |
| 16S-23S rRNA IGS | F | Outer  Forward | GTATGTTTAGTGAGGGGGGTG | 55 | 30 | Variable | (Bunikis et al. 2004) |
|  | R | Outer  Reverse | GGATCATAGCTCAGGTGGTTAG |  |  |  |  |
|  | Fn | Inner  Forward | AGGGGGGTGAAGTCGTAACAAG | 55 | 30 | Variable |  |
|  | Rn | Inner  Reverse | GTCTGATAAACCTGAGGTCGGA |  |  |  |  |
| *flaB* | FlaLL | Outer  Forward | ACATATTCAGATGCAGACAGAGG | 52 | 30 | ~665 | (Barbour et al. 1996) |
|  | FlaRL | Outer  Reverse | GCAATCATAGCCATTGCAGATTGT |  |  |  |  |
|  | 442f | Inner  Forward | GCTGAAGAGCTTGGAATGCAACC | 55 | 30 | ~524 | This paper |
|  | FlaRL | Inner  Reverse | GCAATCATAGCCATTGCAGATTGT |  |  |  | (Barbour et al. 1996) |

*Note: The 16S rRNA screening primers (1A/1B) and the longer region 16S primers (Bf1_24_/Br1_24_) utilized a touchdown PCR approach. The annealing temperature was dropped one degree C in each of the first 10 cycles, followed by 40 additional cycles with annealing temperature as shown in the table.

PCR assays used Promega GoTaq Green master mix (Promega, Madison, WI, USA). Primers were used at 0.5 mM concentration. All PCRs began with an initial denaturation step at 94°C, 2 minutes. Thereafter, cycles consisted of 94°C, 30 sec; annealing temp as indicated in the table for 30 sec; and extension at 72°C, 30 sec for 16S 1A/1B and *flaB* primers or 72°C, 1 min for 16S Bf1_24_/Br1_24_ and IGS primers.

**Table S2**. PCR positivity for *Borrelia* infections summarized across *Pteropus alecto* roosts and sampling sessions. Bats are considered infected if testing positive for at least one of our three markers (i.e., 16S rRNA gene, *flaB* gene, and 16S–23S rRNA ITS).

| **Site** | **Sampling session** | **Session prevalence** | **Total sampled** | **Site prevalence** |
| --- | --- | --- | --- | --- |
| Gympie | 31 January 2019 | .20 | 20 | .20 (4/20) |
| Hervey Bay | 15 July 2018 | 0 | 6 | 0 |
|  | 28 July 2020 | 0 | 30 |  |
| Maclean | 9 July 2018 | 0 | 1 | 0 |
| Mount Ommaney | 17 January 2019 | 0 | 7 | 0 |
| Redcliffe | 25 May 2018 | .04 | 24 | .03 (10/375) |
|  | 27 July 2018 | 0 | 10 |  |
|  | 14 September 2018 | 0 | 19 |  |
|  | 14 December 2018 | .05 | 21 |  |
|  | 8 March 2019 | 0 | 59 |  |
|  | 28 May 2019 | .03 | 30 |  |
|  | 9 July 2019 | .03 | 32 |  |
|  | 10 September 2019 | .03 | 30 |  |
|  | 3 December 2019 | .03 | 32 |  |
| Redcliffe | 3 March 2020 | .07 | 30 | .03 (10/375) |
|  | 11 May 2020 | .07 | 30 |  |
|  | 7 July 2020 | 0 | 30 |  |
|  | 7 September 2020 | 0 | 28 |  |
| **Site** | **Sampling session** | **Session prevalence** | **Total sampled** | **Site prevalence** |
| Toowoomba | 3 June 2018 | .05 | 21 | .007 (3/402) |
|  | 21 July 2018 | .04 | 26 |  |
|  | 8 September 2018 | 0 | 21 |  |
|  | 8 December 2018 | 0 | 22 |  |
|  | 11 January 2019 | 0 | 9 |  |
|  | 15 March 2019 | .02 | 58 |  |
|  | 14 May 2019 | 0 | 30 |  |
|  | 2 July 2019 | 0 | 29 |  |
|  | 23 July 2019 | 0 | 6 |  |
|  | 3 September 2019 | 0 | 30 |  |
|  | 10 December 2019 | 0 | 30 |  |
|  | 10 March 2020 | 0 | 29 |  |
|  | 4 May 2020 | 0 | 30 |  |
|  | 14 July 2020 | 0 | 30 |  |
|  | 1 September 2020 | 0 | 30 |  |

**Figure S1**. Maximum likelihood phylogenetic tree displaying evolutionary relationships between *Borrelia* spp. using the *flaB* gene. This tree was constructed using RAxML 8 (Stamatakis 2014) and a GTR+I+G nucleotide substitution model. Branch support was calculated with 1000 rapid bootstrap replicates.

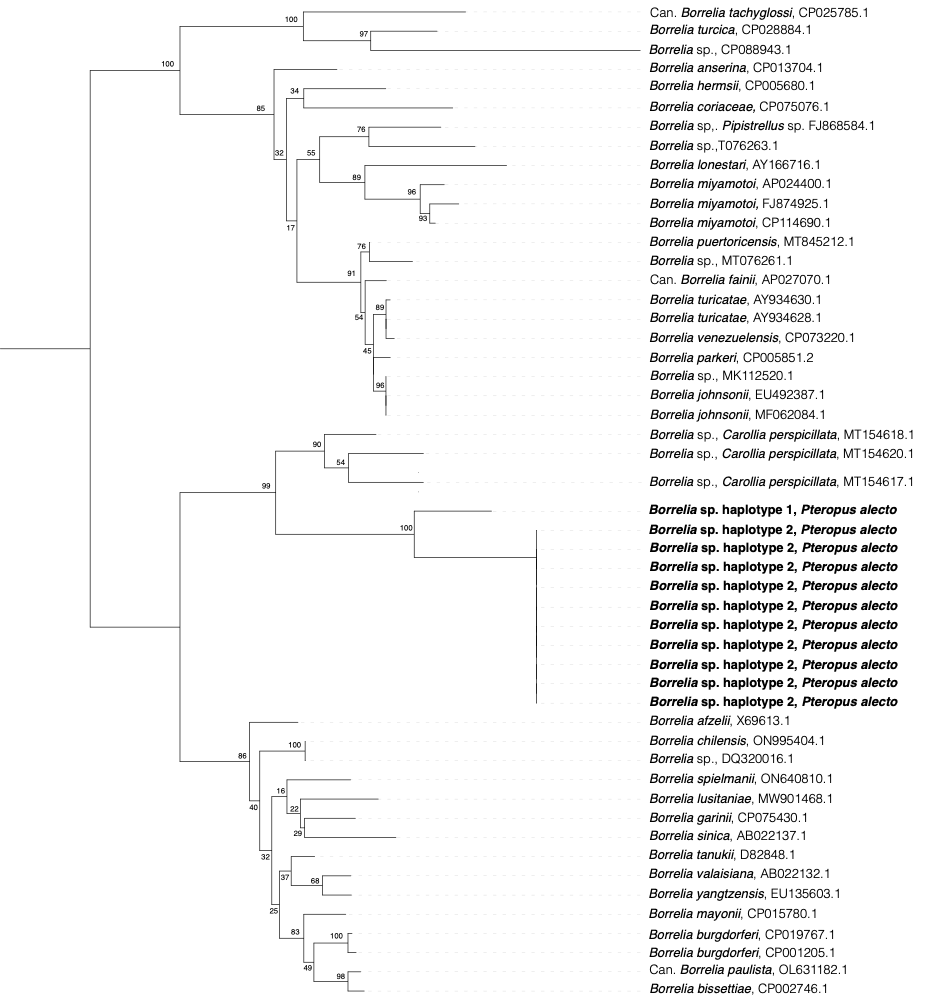

**Figure S2**. Maximum likelihood phylogenetic tree displaying evolutionary relationships between *Borrelia* spp. using the 16S gene. This tree was constructed using RAxML 8 (Stamatakis 2014) and a GTR+I+G nucleotide substitution model. Branch support was calculated with 1000 rapid bootstrap replicates.
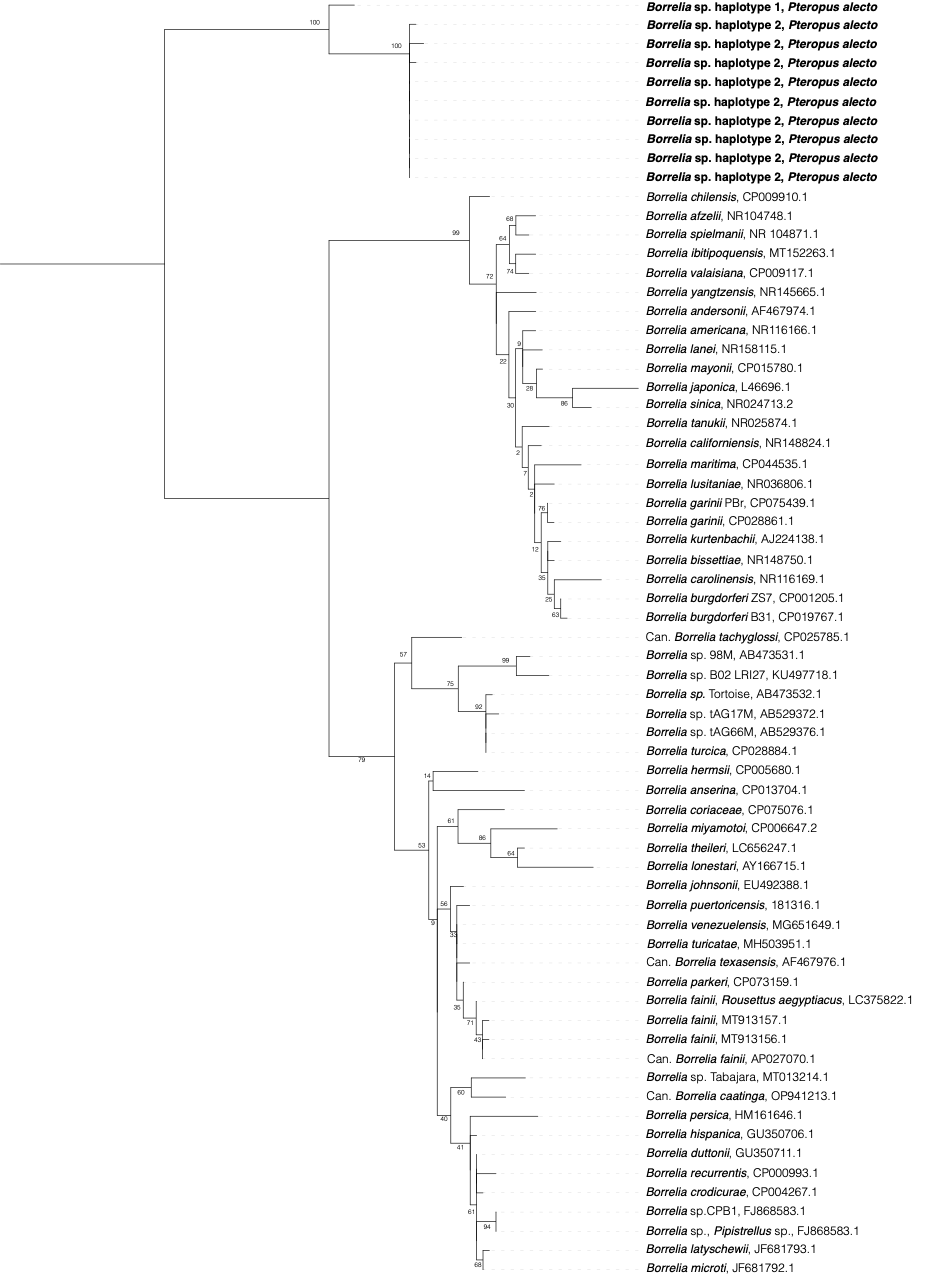
